## Supplemental data for "Resident and recruited macrophages differentially contribute to cardiac healing after myocardial ischemia"

**First Author: Tobias Weinberger**

**Corresponding Author: Christian Schulz**

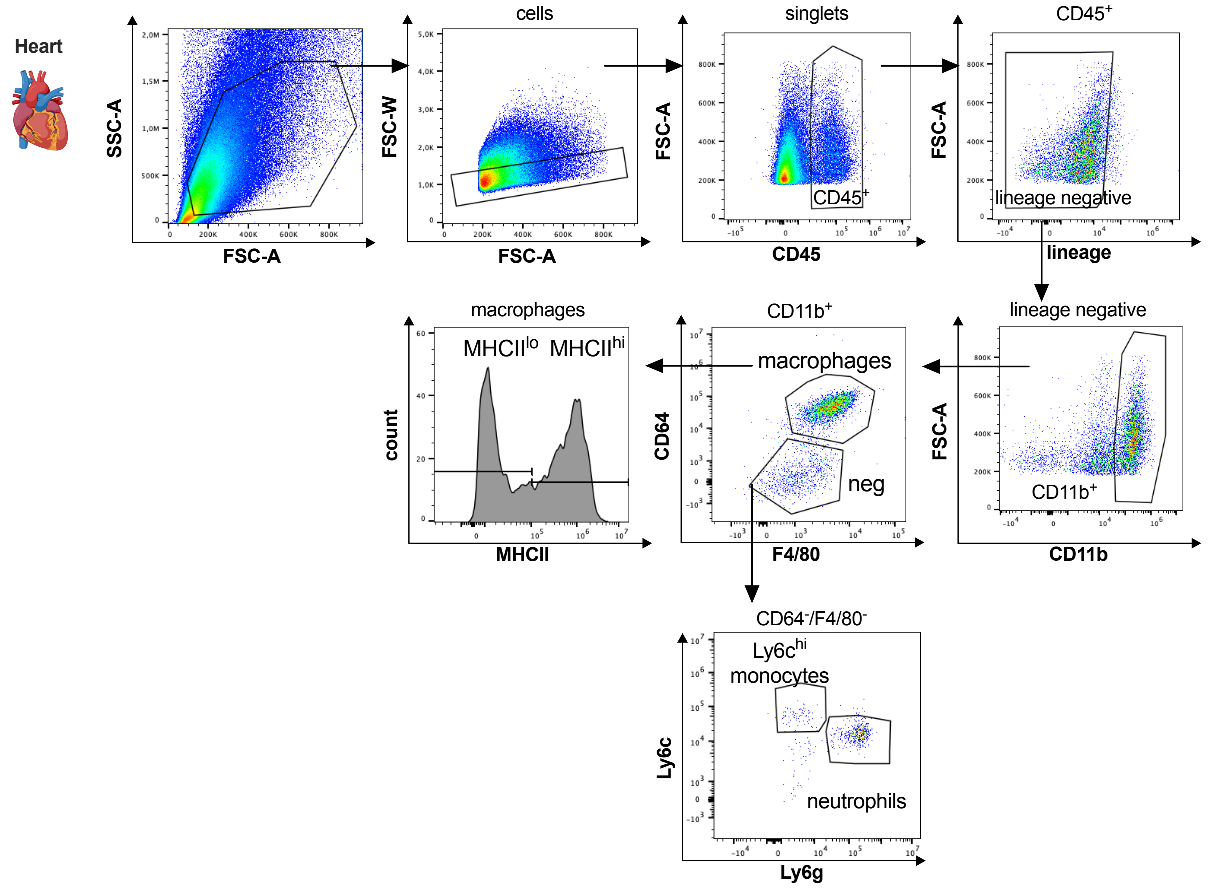


**Fig. S1. Gating strategy for cardiac myeloid immune cells.**

Cardiac myeloid immune cells are gated as single, CD45$^{+}$, lin$^{-}$ (CD11c, Ter119, Tcrß, Nk1.1), CD11b$^{+}$ cells. Macrophages are further characterized as CD64$^{+}$, F4/80$^{+}$ cells with either high or low expression of MHCII, neutrophils as CD64$^{-}$, F4/80$^{-}$, Ly6g$^{+}$ and Ly6c$^{hi}$ monocytes as CD64$^{-}$, F4/80$^{-}$, Ly6g$^{-}$, Ly6c$^{hi}$.

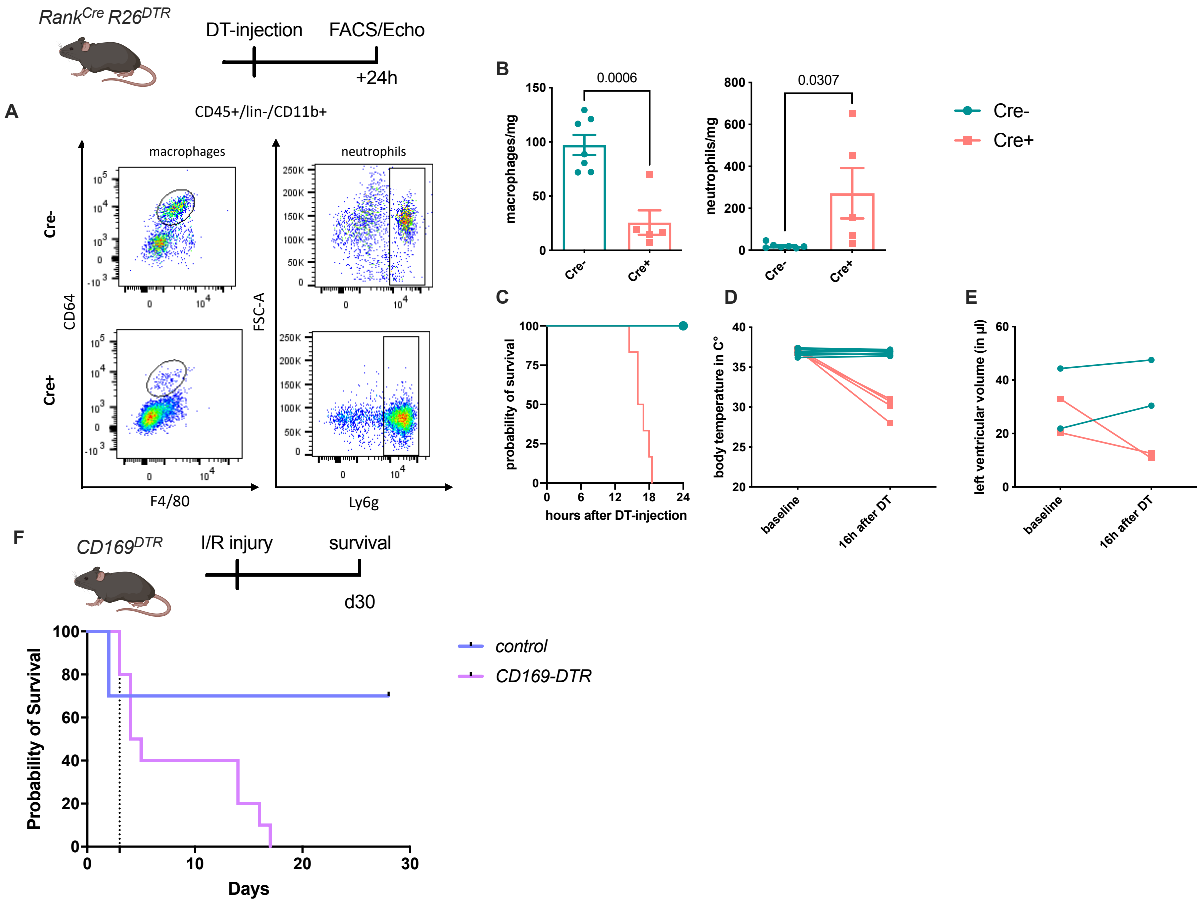


**Fig. S2. DT-mediated depletion of EMP-derived macrophages in *Rank^Cre^Rosa26^DTR^* mice.**

EMP-derived macrophages were depleted using a single DT-injection and analyzed after reaching the termination criteria determined in the ethical regulations (e.g. activity, body score). (A and B) Macrophages and neutrophils were gated as single, CD45$^{+}$, lin$^{-}$, CD11b$^{+}$ and CD64$^{+}$/F4/80$^{+}$ or Ly6g$^{+}$ cells respectively. (C) Survival curve of Cre$^{-}$ (n=4) and Cre$^{+}$ (n=6) mice after DT-injection. (D) Body temperature and (E) left ventricular diastolic volume at baseline and 16h after DT-injection in Cre$^{-}$ and Cre$^{+}$ mice.

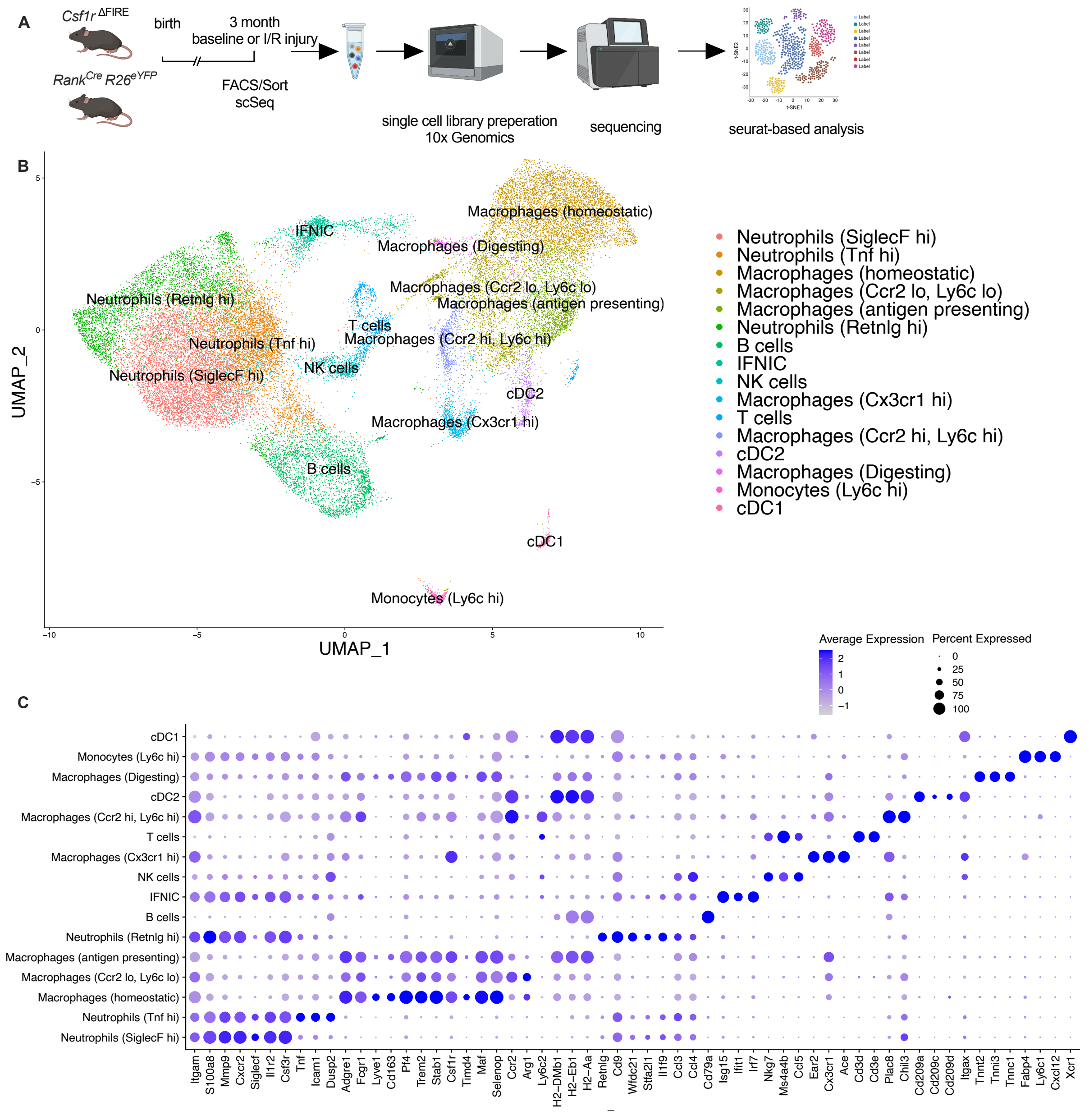


**Fig. S3. Clustering of cardiac immune cells in single cell RNA-analysis.**

**(A)** The whole cardiac immune cell population (single/live/CD45$^{+}$ cells) of *Csf1r*$^{\Delta FIRE}$ and *Rank*$^{Cre}$*Rosa26*$^{eYFP}$ mice were FACS-sorted and single cell library preparation was performed on the 10x platform (n=3 for *Csf1r*$^{\Delta FIRE/+}$ and *Csf1r*$^{\Delta FIRE/\Delta FIRE}$ in baseline conditions, n=1 for *Rank*$^{Cre}$*Rosa26*$^{eYFP}$ in baseline conditions; n=3 for *Csf1r*$^{\Delta FIRE/+}$ and *Csf1r*$^{\Delta FIRE/\Delta FIRE}$ 2 days after I/R-injury; in mice subjected to I/R injury macrophages were harvested from the ischemic tissue). **(B)** In total 35759 cells(13148 cells in baseline and 22611 cells after I/R injury d2) were analyzed. Using Seurat-based clustering we identified 16 different immune clusters. Marker genes for each individual cluster are depicted in **(C)**.

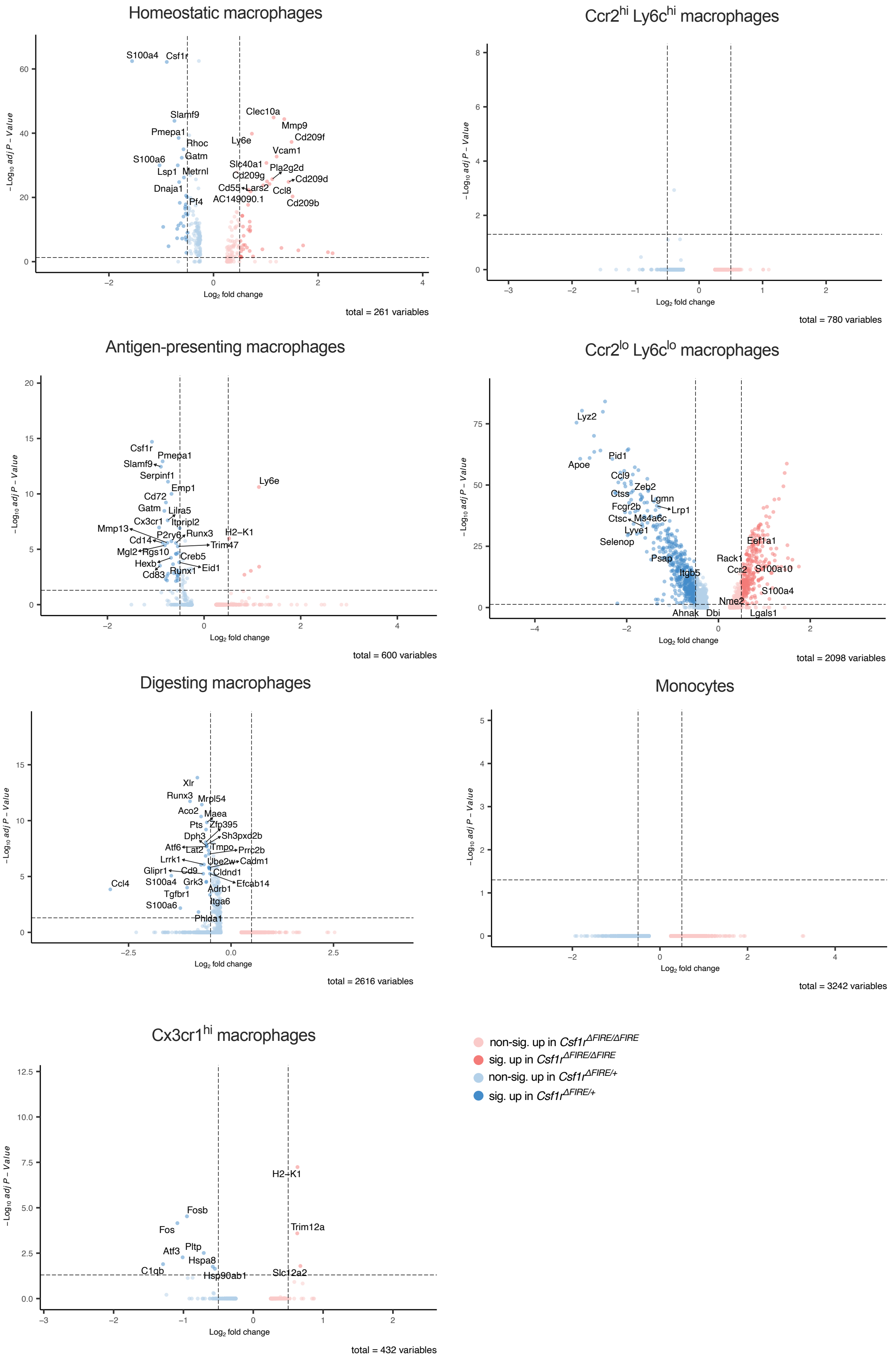


**Fig. S4. Differential gene expression in monocyte and macrophage clusters in baseline conditions in *control* and *ΔFIRE* mice.**

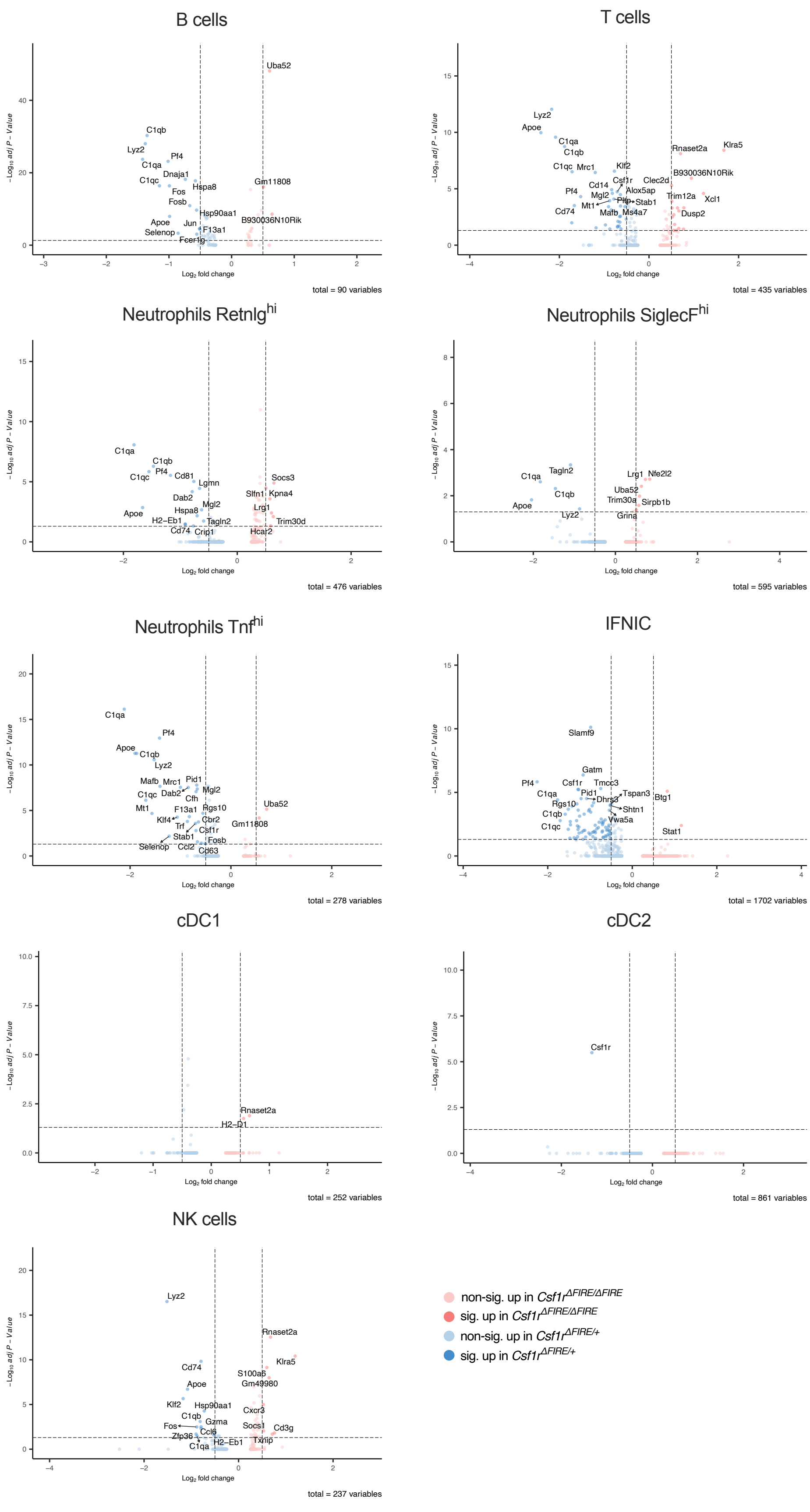


**Fig. S5. Differential gene expression in non-macrophage clusters in baseline conditions in *control* and *ΔFIRE* mice.**

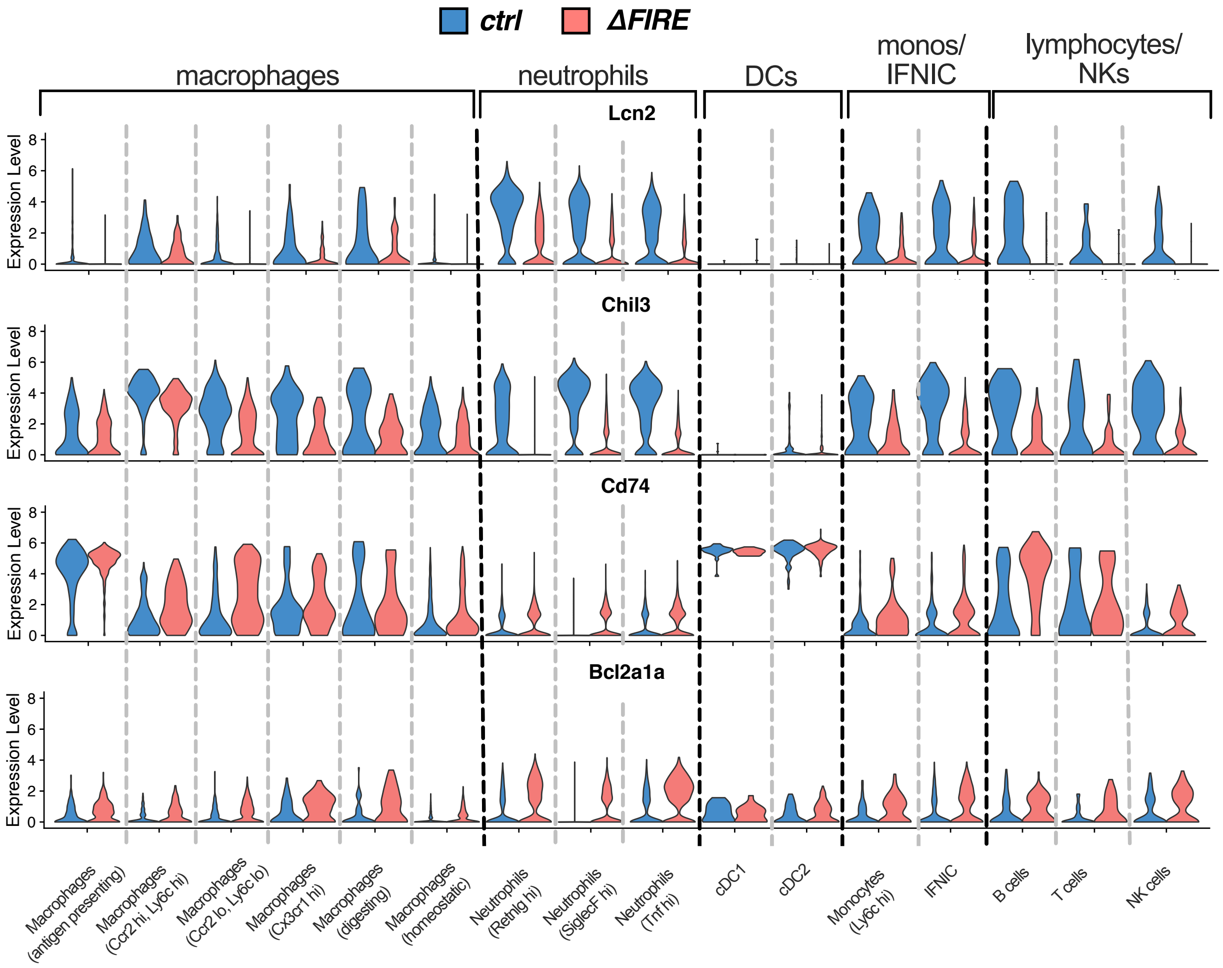


**Fig. S6. Violin plots comparing expression of Lcn2, Chil3, CD74 and Bcl2a1a in the different immune cell clusters after I/R in control and ΔFIRE mice..**

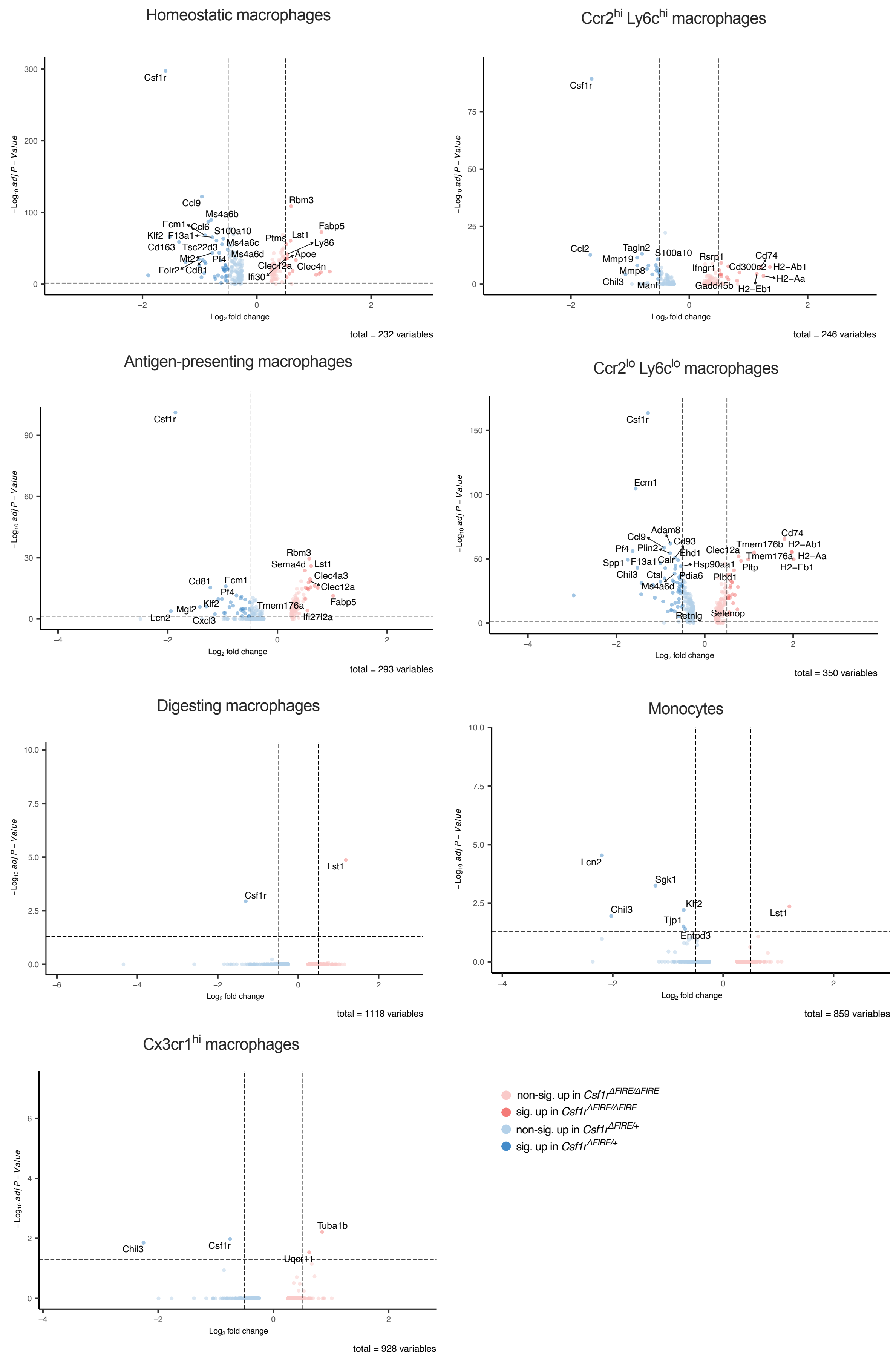


**Fig. S7. Differential gene expression in monocyte and macrophage clusters after I/R in *control* and *ΔFIRE* mice.**

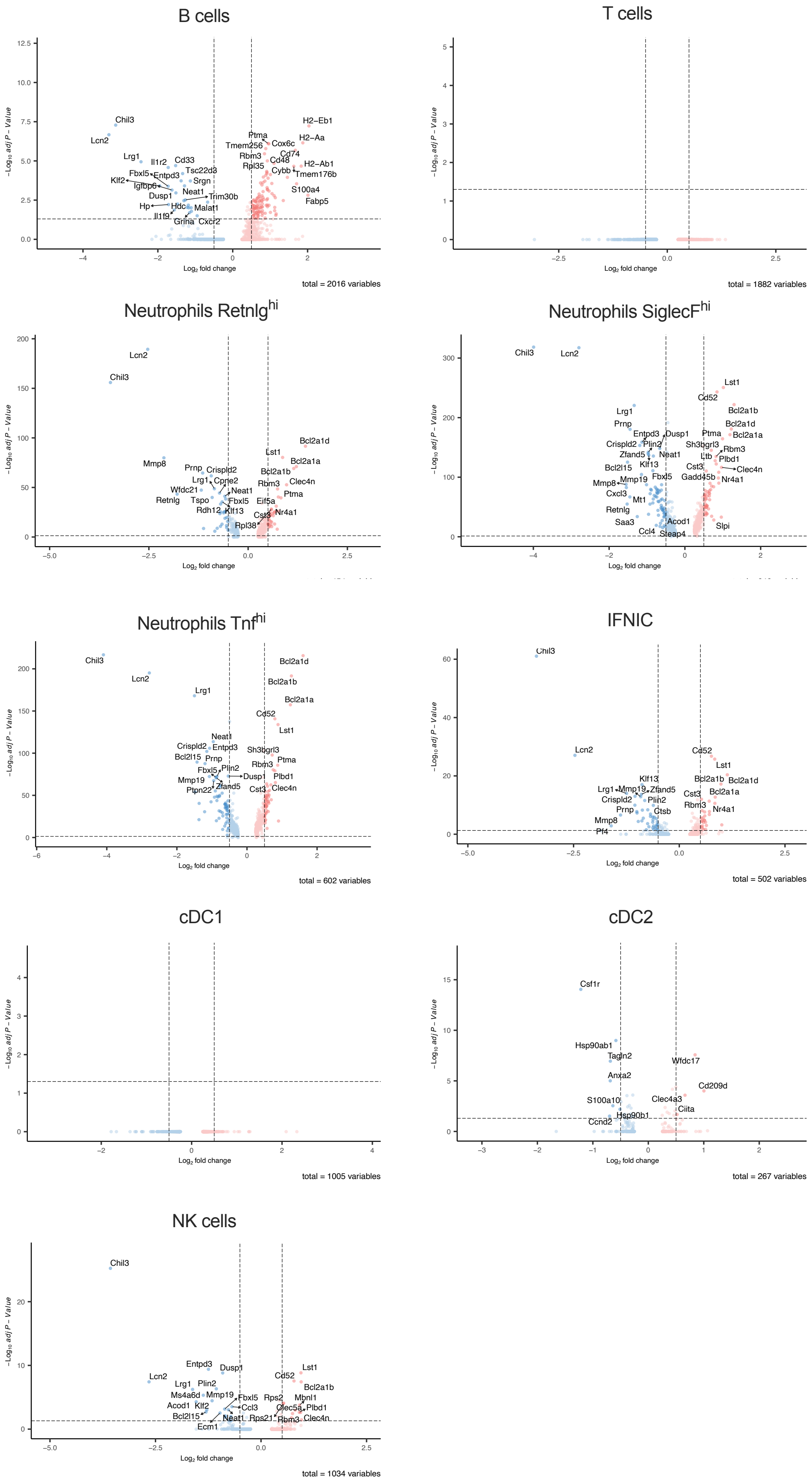


**Fig. S8. Differential gene expression in non-macrophage clusters after I/R in in *control* and *ΔFIRE* mice.**

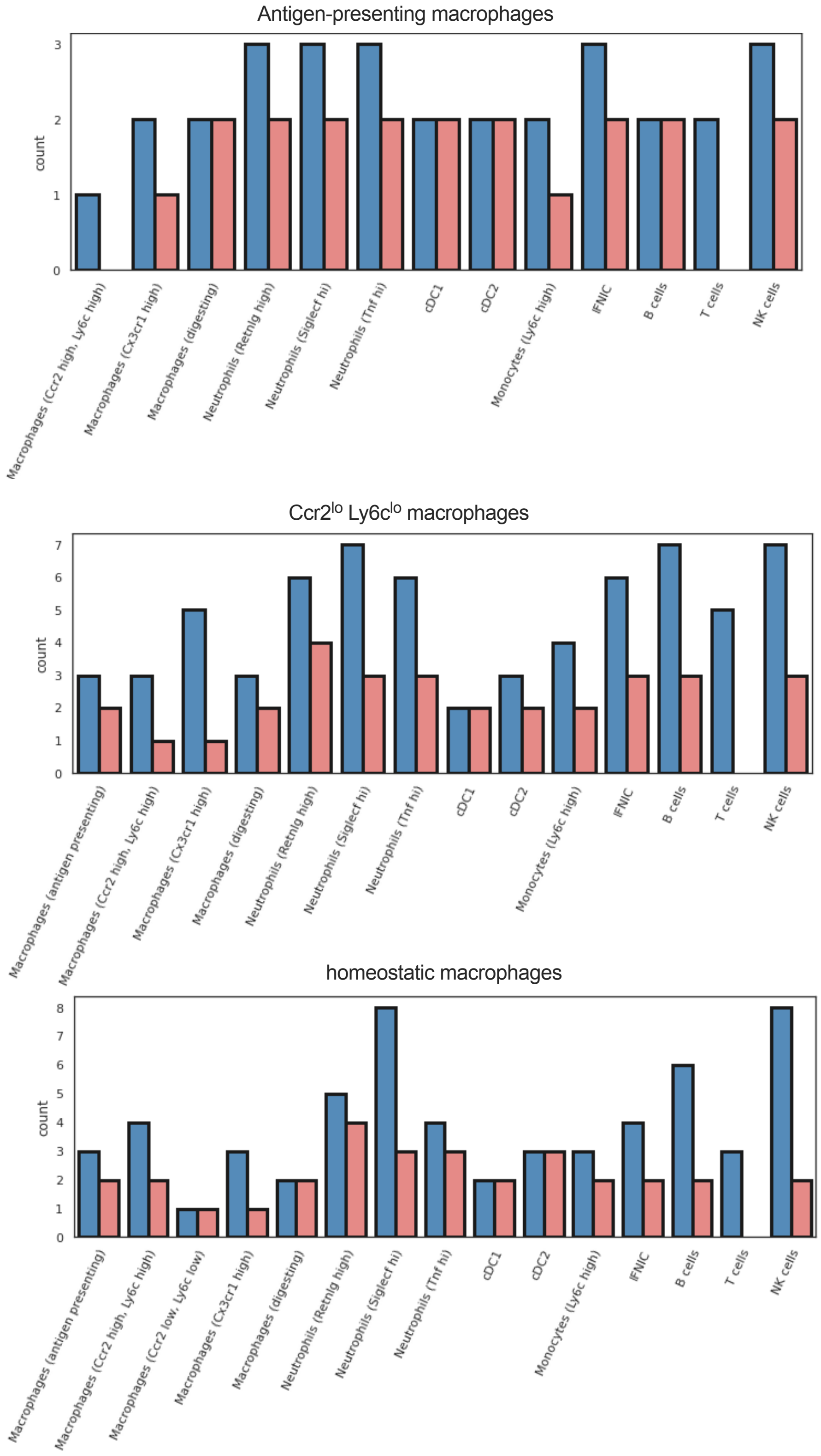


**Fig. S9. Differences in outgoing signals in macrophage subpopulations in I/R injury in *Csf1r*$^{\Delta FIRE/+}$ and *Csf1r*$^{\Delta FIRE/\Delta FIRE}$ mice.**

Number of cell-cell interactions (with communication score > 6) outgoing from (a) antigen-presenting macrophages , (b) *Ccr2*$^{lo}$*Ly6c*$^{lo}$ macrophages and (c) homeostatic macrophages to other immune cell clusters in *Csf1r*$^{\Delta FIRE/+}$ (blue) and *Csf1r*$^{\Delta FIRE/\Delta FIRE}$ (red).

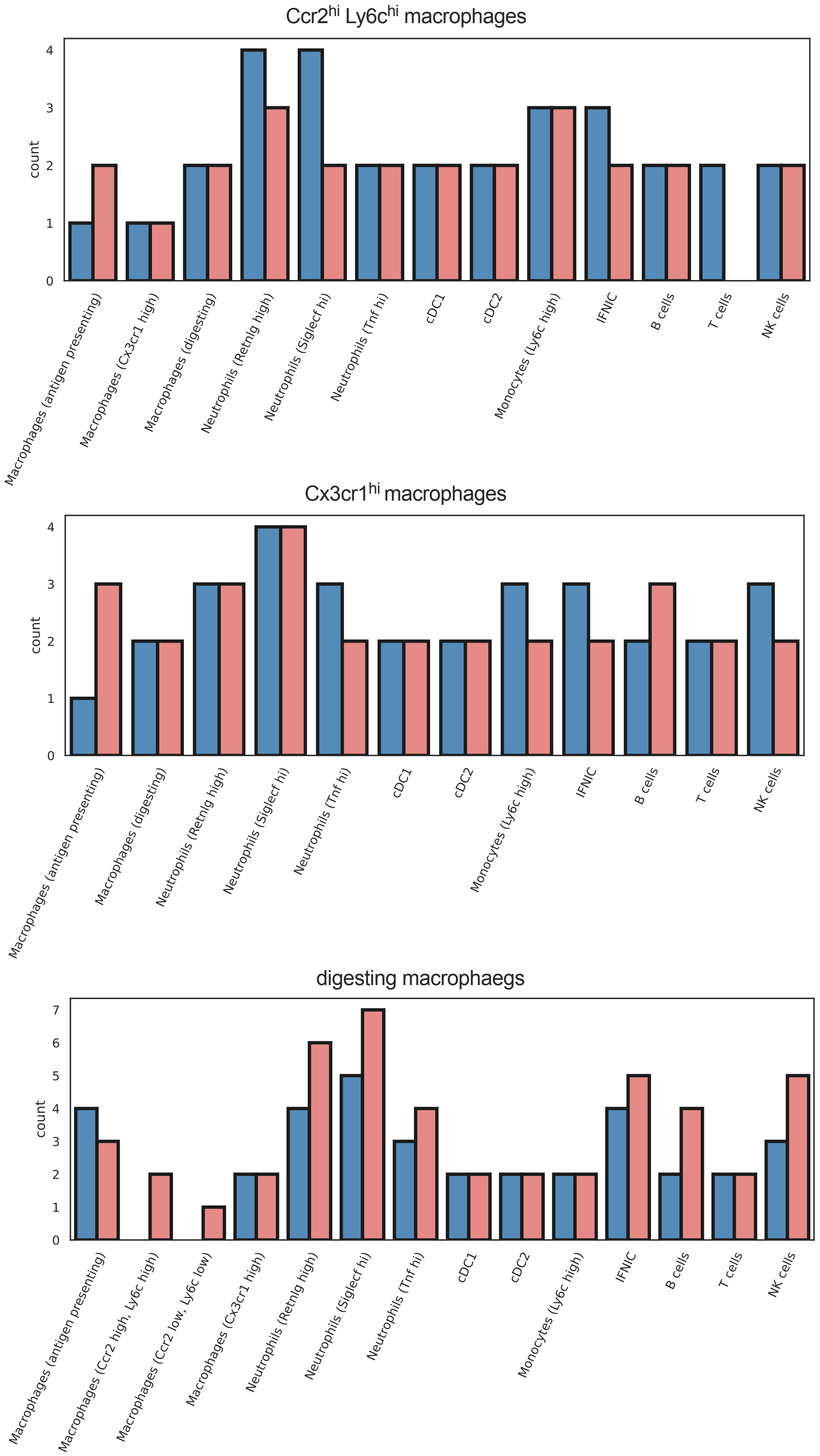


**Fig. S10. Similar outgoing signals in different macrophage subpopulations in I/R injury in *Csf1r*$^{\Delta FIRE/+}$ and *Csf1r*$^{\Delta FIRE/\Delta FIRE}$ mice.**

Number of cell-cell interactions (with communication score > 6) outgoing from (a) Ccr2 hi, Ly6c high macrophages, (b) Cx3cr1 high macrophages (c) digesting macrophages to other immune cell clusters in *Csf1r*$^{\Delta FIRE/+}$ (blue) and *Csf1r*$^{\Delta FIRE/\Delta FIRE}$ (red).

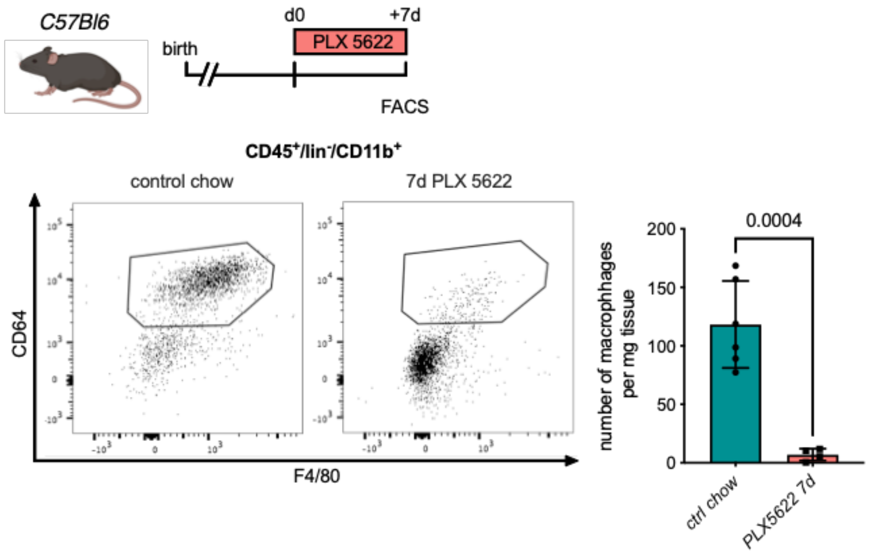


**Fig. S11. Macrophage depletion using the Csf1r-inhibitor PLX5622.**

Flow cytometry analysis of hearts from *C57Bl6* mice treated with PLX5622 for 7 days, (left) representative flow cytometry showing cardiac macrophages (CD45+/lin-/CD11b+/F4/80+/CD64+ cells) and (right) number of cardiac macrophages in animals fed control chow or PLX5622 (n=6 for control chow and n=4 for PLX5622, each individual experiments).

| Antibody | Dilution | Clone | Fluorescence | Company | Catalog number |
|---|---|---|---|---|---|
| **Flow Cytometry** | | | | | |
| **CD115** | 1:100 | AFS98 | BV421 | Biolegend | 135513 |
| **CD115** | 1:100 | AFS98 | APC | Biolegend | 135509 |
| **CD11b** | 1:100 | M1/70 | PE-Cy7 | BD Biosciences | 552850 |
| **CD11b** | 1:100 | | APC-Cy7 | | |
| **CD11c** | 1:100 | N418 | PE | Biolegend | 117308 |
| **CD11c** | 1:100 | | Fitc | | |
| **CD16/CD32 (FcγRIII/FcγRII)** | 1:100 | 2.4G2 | uncoupled | BD Biosciences | 553142 |
| **CD19** | 1:200 | | | | |
| **CD64** | 1:100 | X54-5/7.1 | APC | Biolegend | 139306 |
| **CD45** | 1:100 | | BUV395 | | |
| **CD45** | 1:100 | 30-F11 | PerCP | Biolegend | 103130 |
| **CD45.1** | 1:100 | A20 | Fitc | BD Biosciences | 110706 |
| **CD45.2** | 1:100 | 104 | APC-Cy7 | BD Biosciences | 560694 |
| **F4/80** | 1:100 | BM8 | BV421 | Biolegend | 123131 |
| **Gr-1** | 1:100 | RB6-8C5 | bio | eBio | 13-5931-82 |
| **Gr-1** | 1:100 | | | | |
| **Ly6G** | 1:100 | 1A8 | BV605 | Biolegend | 127639 |
| **NK1.1** | 1:100 | PK136 | PE | Biolegend | 108708 |
| **NK1.1** | 1:100 | | Fitc | | |
| **TCR-ß** | 1:100 | 1B3.3 | PE | Biolegend | 109208 |
| **TCR-ß** | 1:100 | | Fitc | | |
| **Ter119** | 1:100 | TER-119 | PE | Biolegend | 116208 |
| **Ter119** | 1:100 | | Fitc | | |
| **I-A/I-E Antibody** | 1:100 | M5/114.15.2 | PE/Cy7 | Biolegend | 107630 |
| **SYTOX™ Orange** | 1:1000 | | PE | Thermofisher | S11368 |
| **CD45 MicroBeads, mouse** | 1:100 | 30F11.1 | | Miltenyi Biotec | 130-052-301 |
| **Immunohistology** | | | | | |
| **anti-GFP** | 1:100 | Polyclonal, Rabbit | uncoupled | Invitrogen | A-11122 |
| **CD68** | 1:200 | FA11 | uncoupled | BioRad | MCA1957 |
| **Wheat Germ Agglutinin** | 1:100 | | Alexa Fluor 350 | ThermoFisher Scientific | W11263 |
| **AF488 goat anti-rabbit** | 1:200 | Polyclonal | Alexa Fluor 488 | Invitrogen | A-11034 |
| **AF 555 goat anti rat** | 1:200 | Polyclonal | Alexa Fluor 555 | Invitrogen | A-21343 |

**Table S1. Antibody list.**
